# Intrinsic Units: Identifying a system’s causal grain

**DOI:** 10.1101/2024.04.12.589163

**Authors:** William Marshall, Graham Findlay, Larissa Albantakis, Giulio Tononi

**Author notes:** These authors contributed equally to this work (William Marshall, Graham Findlay).

## Abstract

Integrated information theory (IIT) aims to account for the quality and quantity of consciousness in physical terms. According to IIT, a substrate of consciousness must be a system of units that is a maximum of intrinsic, irreducible cause-effect power, quantified by integrated information (*φ*_*s*_). Moreover, the grain of each unit must be the one— from micro (finer) to macro (coarser)—that maximizes the system’s intrinsic irreducibility (i.e., maximizes *φ*_*s*_). The units that maximize *φ*_*s*_ are called the intrinsic units of the system. This work extends the mathematical framework of IIT 4.0 to assess cause-effect power at different grains and thereby determine a system’s intrinsic units. Using simple, simulated systems, we show that the cause-effect power of a system of macro units can be higher than the cause-effect power of the corresponding micro units. Two examples highlight specific kinds of macro units, and how each kind can increase cause-effect power. The implications of the framework are discussed in the broader context of IIT, including how it provides a foundation for tests and inferences about consciousness.

## 1 Introduction

One goal of the scientific study of consciousness is to ascertain its neural substrate. Much attention has been given to the question of which regions of the brain support consciousness [9, 22]. No less important, but less often considered, is the question of the units constituting the substrate of consciousness and their “grain.” Are the units individual neurons, synapses, groups of neurons, or the smallest units that we can possibly manipulate and observe? Is a unit’s state over a hundred milliseconds, or one millisecond, or one second what matters for consciousness? These issues are not only empirical, but call for a theoretical understanding of why certain brain regions qualify as a substrate of consciousness, while others do not, and why the grain of each unit within a substrate is what it is.

Integrated information theory (IIT) aims to account for consciousness—its quality and quantity—by starting from phenomenology and identifying its essential properties—the *axioms* of phenomenal existence—that are true of every conceivable experience: *existence, intrinsicality, information, integration, exclusion*, and *composition* [1].

The axioms of phenomenal existence are formulated as corresponding physical properties, called *postulates*, that must be satisfied by the substrate of consciousness. Physical existence is defined operationally in terms of cause-effect power, and the postulates therefore require that the substrate of consciousness has cause-effect power upon itself (intrinsicality), in a way that is specific (information), unitary (integration), definite (exclusion), and structured (composition). In principle, by evaluating whether and in what way a candidate substrate satisfies all of the postulates, one can evaluate whether and in what way it is conscious, with no additional ingredients.

IIT provides a precise mathematical definition of intrinsic, specific, unitary, definite, and structured cause-effect power, so that it is possible (in principle) to assess whether any given physical system satisfies the postulates. IIT’s framework has been refined over time [27, 6, 23, 1], including several recent developments [14, 7, 19] (for applications outside of consciousness science see [2, 20, 3]). The current framework—IIT 4.0—aims to provide a complete, self-consistent formulation of the postulates in mathematical terms, guided by clearly articulated methodological and ontological principles [1].

According to IIT, a substrate of consciousness, called a *complex*, is a set of units in a state whose intrinsic cause-effect power is maximally irreducible, as measured by its *integrated information* (*φ*_*s*_). In turn, the grain of each unit is the one that maximizes the complex’s *φ*_*s*_. Initial work on determining a complex’s intrinsic units introduced the notion of a *macro unit* —a coarser unit derived from a set of finer units—and demonstrated that the cause-effect power of a system, as measured by effective information [29], could peak when constituted of macro units [16]. Further studies explored how and why systems of macro units can have higher cause-effect power than the systems of micro units upon which they supervene [15, 18, 12].

The goal of the current work is to provide a mathematical framework for identifying a system’s *intrinsic units*— those that constitute it from its intrinsic perspective—based on IIT 4.0 [1], in a way that incorporates the theory’s postulates explicitly. In Section 2, we review IIT 4.0’s mathematical framework for measuring the cause-effect power of systems of micro units, and then extend this framework to systems containing macro units. In Section 3, the updated framework is applied to simple systems, demonstrating that macro-grain systems can have higher cause-effect power than the micro-grain systems on which they supervene. In Section 4, we provide a brief discussion of the importance of this framework for future work.

## 2 Theory

The IIT framework provides both the means to identify substrates of consciousness (by computing integrated information) and to account for “what it is like” to be that substrate (by *unfolding* its cause-effect structure). These procedures are described in detail elsewhere [19, 1]. Unfolding, though a crucial aspect of IIT, is not necessary for expanding IIT’s framework to consider cause-effect power at macro grains. This is because the unfolding process is the same for all substrates, regardless of grain. For this reason, we will not discuss unfolding here, and refer interested readers to [1].

In this section, we first highlight features of IIT’s mathematical framework that are necessary for its extension to macro grains. Next, we introduce a definition for macro units constituted of micro units, and extend the formalization of IIT’s postulates to systems containing macro units, by introducing the notion of intrinsic units. Finally, we extend the mathematical framework so it can be used to measure the cause-effect power of systems containing macro units.

### 2.1 Cause-effect power at the micro grain

According to IIT, something can be said to exist physically if it can “take and make a difference.” (i.e., bear a cause and produce an effect). Operationally, it must be possible to manipulate the system’s units (change their state) and observe the result.

The starting point of IIT’s mathematical framework is a stochastic model for a physical universe *U* = {*U*_1_, …, *U*_*n*_} of *n* interacting units with state space Ω_*U*_ = {0, 1}^*n*^. We define *u* as the set of units *U* in a particular state. More precisely, *u* = { (*U*_*i*_, state(*U*_*i*_)) : *U*_*i*_ ∈*U*} is a set of tuples, where each tuple contains a unit and the state of that unit. This construction allows us to define set operations over *u* that consider both the units and their states. We further denote Ω_*U*_ to be the set of all such tuple sets, corresponding to all the possible states of *U*. Because the physical existence of *U* is formulated operationally as cause-effect power, *U* is defined by its potential interactions, assessed in terms of conditional probabilities. We denote the complete *transition probability matrix* (TPM) of a universe *U* over a system update u *→* ū as:

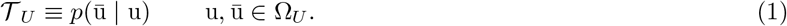

The TPM provides a complete description of *U* at the micro grain—the finest grain at which manipulation and observation is possible, with nothing omitted from the causal model—which means that we can determine the conditional probabilities in (1) for every system state, with *p*(ū| u) = *p*(ū |do(u)) [3, 17, 5, 24] (where the “do-operator” do(u) indicates that u is imposed by intervention). This implies that *U* corresponds to a complete causal network [3].

According to IIT, 𝒯_*U*_ does not merely *describe* the physical universe, but rather *is* the physical universe, because there is no need for intrinsic properties (things like mass, charge, spin, etc.) that would “underlie” or come prior to *𝒯* _*U*_ —there is just cause-effect power. What a micro physical system is, is just its TPM, and vice versa [1, 11].

Because *U* is assumed to be a complete causal network, it will not exhibit “instantaneous causation.” More formally, the individual random variables *U*_*i*_ ∈ *U*, conditional on the preceding state of *U*, are independent from each other:

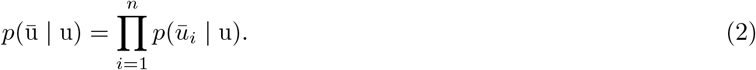

For an extension of IIT to quantum systems, where model completeness may not imply such conditional independence, see [4].

*Micro units* are the finest units that can be observed and manipulated, satisfying the minimal requirements for cause-effect power (i.e., physical existence) [1]. Accordingly, micro units must have exactly two states—”this way” and “not this way.” One state is simply the complement of the other, whichever state is picked, with no further qualification. Having more than two states would imply an internal mechanism distinguishing among “this way”, “that way”, and “the other way,” that would remain hidden within each unit, inaccessible to manipulation or observation.

For any *candidate substrate* (also called a *candidate system*) *S* ⊆ *U* in a state *s* ⊆ *u*, the IIT 4.0 framework defines its *system integrated information φ*_*s*_(*s*) [19, 1]. Based on the postulates of intrinsicality, information, and integration, *φ*_*s*_(*s*) quantifies how the system specifies a *cause-effect state* as a whole, above and beyond how it specifies the same cause-effect state as independent parts [1]. Per the *principle of minimal existence*, which states that “nothing exists more than the least it exists” (e.g., “a chain is only as strong as its weakest link”), the comparison between the whole and its parts is performed by partitioning the system and evaluating the impact of the *minimum partition*—the partition over which the system is least irreducible [1]. The system integrated information, *φ*_*s*_, is defined as the *intrinsic information* of the whole [8, 7], relative to the parts specified by its minimum partition. We do not present the full definition or algorithm for obtaining *φ*_*s*_ here—only the parts that are relevant for extending the framework to macro units.

For any candidate system, *φ*_*s*_(*s*) is defined based on two system-specific transition probability matrices, *𝒯*_*c*_ and *𝒯*_*e*_ (for describing causes and effects respectively). The system TPMs are computed by *causally marginalizing* the units *W* = *U \ S* conditional on the current state *u*, as described below. *W* in state *w* are referred to as the system’s *background units* or *background conditions*, and they may partly enable its cause-effect power [1]. *𝒯*_*c*_ and *𝒯*_*e*_ therefore capture intrinsic cause-effect power of the system within the context of a set of background conditions.

For evaluating effects of the current state, the state of the background units is fully determined by the current state of the universe (*w* = *u \s*). The corresponding TPM, *𝒯*_*e*_, is used to identify the effect of the current state:

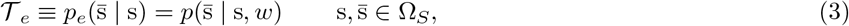

For evaluating causes of the current state, knowledge of the current state is used to compute the probability distribution over possible past states of the background units, which is not necessarily uniform or deterministic. This distribution is computed using Bayes’ rule, assuming a uniform marginal distribution of the previous state:

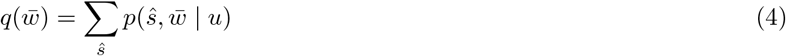

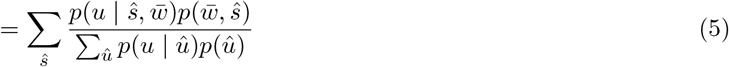

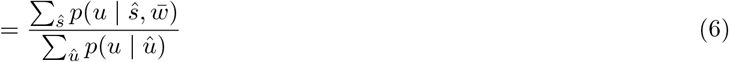

The corresponding TPM, *𝒯* _*c*_, is a weighted average of transitions probabilities (weighted by 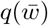) over possible states of background units:

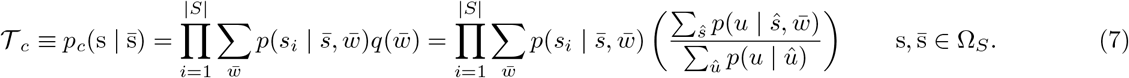

Note that *𝒯*_*c*_ is computed one unit at a time, and then a product is used to compute probabilities for the whole system. This is done to remove correlations among the units due to using a weighted average of background conditions, rather than cause-effect power of the units *per se*.

According to the exclusion postulate, a substrate of consciousness must be definite: there must be a reason why it consists of these units, and not others. The reason is provided by the *principle of maximal existence*, which states that among competing existents, the one that actually exists is the one that exists the most. Furthermore, if maximal existence is the sufficient reason for a complex being supported by a given set of units, it is also the sufficient reason for not being supported by subsets, supersets, or parasets of that set. This implies that complexes cannot overlap, in line with the notion that a micro unit’s cause-effect power should not be counted multiple times [28].

Since existence as *one* entity is quantified by integrated information *φ*_*s*_, complexes can be identified as maxima of *φ*_*s*_[1]. That is, *s* is a complex if:

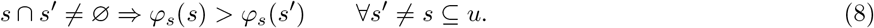

Put simply, we compare a candidate system *s* ⊆ *u* to all other potential candidate systems *s*^*′*^ ⊆ *u*, and ensure that its system integrated information is greater than any subset, superset, or paraset of itself (i.e., any overlapping candidate system). Over a universal substrate *u*, non-overlapping complexes are identified recursively (first-maximal complex, then second-maximal complex, and so on).

Finally, it follows from exclusion and the principle of maximal existence that a complex should not only have greater *φ*_*s*_ than overlapping candidate systems constituted of micro units, but also any overlapping systems constituted of macro units. In the next section, we extend IIT’s mathematical framework to permit evaluation of *φ*_*s*_ for systems containing macro units.

### 2.2 Intrinsic Units

The grain of each unit from a system’s intrinsic perspective (its “intrinsic units”) is the one at which the system’s *φ*_*s*_ is maximized. It is not a requirement that a complex’s units all share the same grain, so a *system’s grain* refers not to a single grain, but rather to its particular configuration of units at their particular (possibly hetereogenous) grains. A system has a macro grain if it contains at least one macro unit. Evaluating *φ*_*s*_ for candidate systems at all possible grains requires extending IIT’s mathematical framework as follows.

#### 2.2.1 Meso and macro units

Starting from a set of micro units within *U*, a macro unit can be obtained by macroing “over units” (when a macro unit has more than one constituent unit), “over updates” (when a macro unit has more than one update step), or both. Previous work referred to macroing as being “over space” and/or “over time” [16, 18], but we avoid these terms here, because of their metaphysical implications. The IIT framework does not require spacetime to be fundamental.

As mentioned above, micro units are the finest units that can be observed and manipulated, satisfying the minimal requirements for cause-effect power (i.e., physical existence) [1]. These “atoms” of cause-effect power cannot be partitioned into finer constituents, their updates cannot be partitioned into finer updates, and they cannot have more than two states—the minimum necessary to bear a cause and produce an effect.

Macro units, from the intrinsic perspective of a complex, are its “units” of cause-effect power and must have a repertoire of exactly two states—the minimum necessary to bear a cause and produce an effect. Having more than two states would bring into play internal mechanisms distinguishing among “this way”, “that way”, and “the other way”, within each unit. These internal mechanisms—provided by the macro unit’s internal micro constituents—would contribute to cause-effect power among macro units while remaining hidden inside them, inaccessible to partitions (resulting in misattribution of causal power from finer grains to coarser ones). Of course, from the extrinsic perspective of an experimenter unconcerned with the separation of grains, non-binary macro states are available for observation and manipulation, and can reveal important causal properties of a substrate (see Section 4).

For the purpose of defining units at different grains, we assume that *U* = {*U*_1_, *U*_2_, …, *U*_*n*_} is a set of micro units. A macro unit *J* constructed from micro units has four aspects:

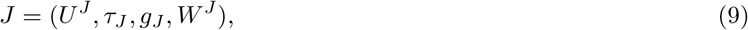

where *U*^*J*^ ⊆ *U* are its *micro constituents, τ*_*J*_ ∈ ℤ^+^ is its *update grain* in terms of micro updates, and *g*_*J*_ a mapping from the states of *U*^*J*^ over a sequence of *τ*_*J*_ micro updates to the state of *J* :

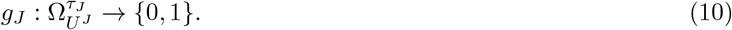

The fourth element of a macro unit, *W*^*J*^, is its *apportionment* of the complex’s background conditions. For a system at a micro update grain, the current state of its background units provides the context that may partly enable the system’s cause-effect power [1]. At a macro update grain, this context is provided not only by the current state of a system’s background units, but also by the way their state changes over multiple micro updates. This means that background units can potentially mediate cause-effect power among units. However, the cause-effect power of units should not be counted multiple times. Accordingly, there cannot be any overlap among the micro units mediating the effects of different macro units. Background units are thus partitioned into disjoint sets and apportioned to specific macro units in a way that maximizes the complex’s *φ*_*s*_. *W*^*J*^ can be ignored at the micro update grain, because a single micro update does not provide any opportunity for background units to mediate interactions among a system’s units.

Constructing macro units directly from micro units is a special case of a more general framework. A macro unit may also be built from constituents *V* ^*J*^ that are themselves macro units at a finer grain—called *meso units*—and the same may be true for the meso units’ constituents, and so forth. That is, a macro unit may be built from one or more levels of meso units sandwiched between it and its constituent micro units *U*^*J*^ (Figure 1). Formally, there is no difference between macro units and meso units, but for clarity we will hereafter reserve the term “macro” for the grain of an intrinsic unit (when it is not a micro unit), and “meso” for any intermediate grains.

**Figure 1:**
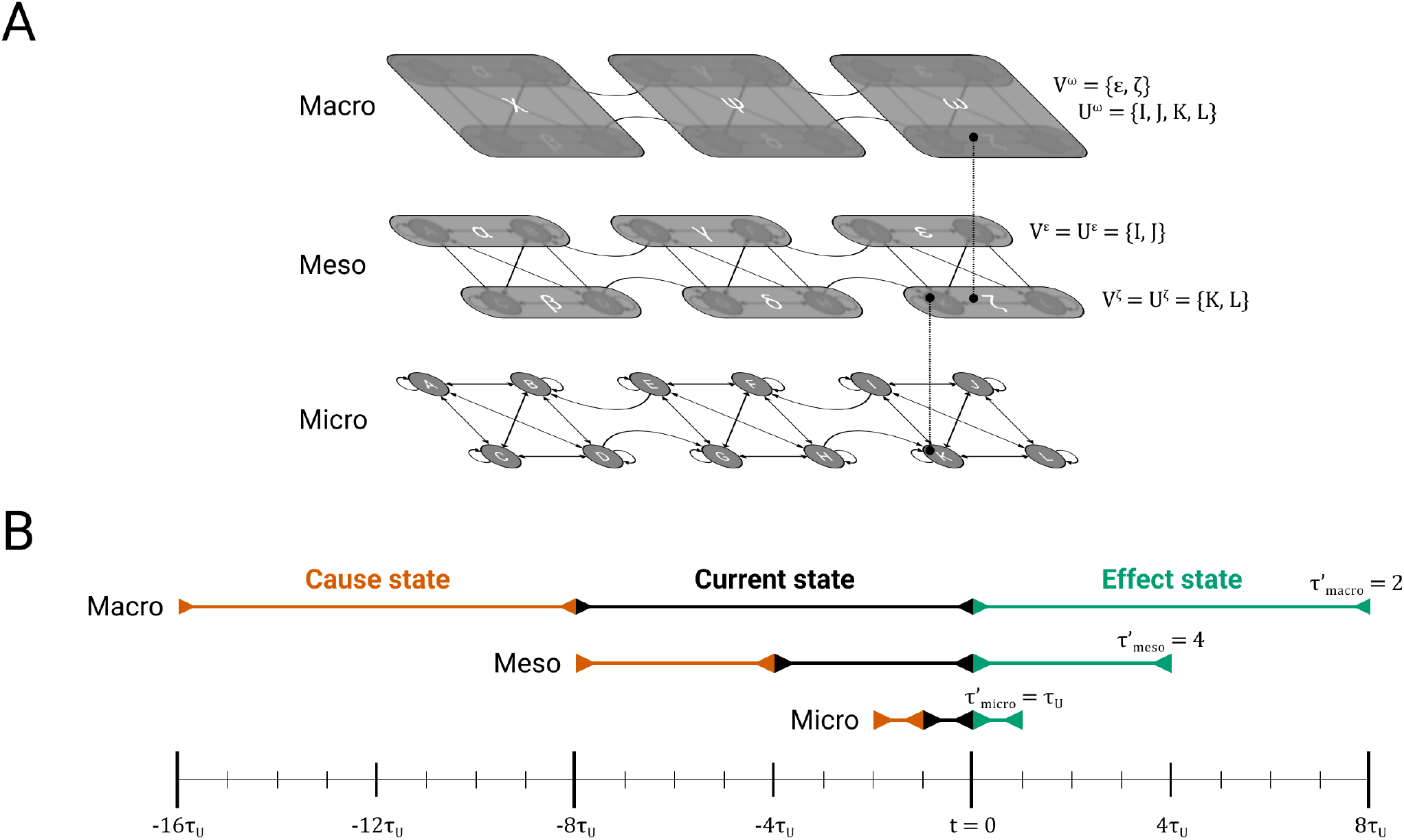
From micro to macro units. (**A**) A universe *U* = {*A, B, C, D, E, F, G, H, I, J, K, L*}, with unspecified transition probability function 𝒯_*U*_. Although some intrinsic cause-effect power may be associated with units at this micro grain, it is also possible that intrinsic cause-effect power is highest at a macro grain. For example, it might be maximal for the macro system {*χ, ψ, ω*}, which would mean that this system exists from its own perspective as a system of three macro units. The framework provided in this paper will allow us to assess if this is the case, including whether a macro unit, say *ω*, can be built using micro constituents *U* ^*ω*^ = {*I, J, K, L*}, possibly with intermediate meso constituents *V* ^*ω*^ = {*ϵ, ζ*}. (**B**) In addition to defining macro states over groups of units, it is also possible to define macro states over updates of *U*. We depict one hypothetical scenario in which macro units have an update grain equal to 2 meso updates 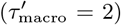, meso updates have an update grain equal to 4 micro updates 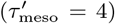, and the micro update 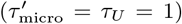 is inherited from *U*. The macro state of a unit is always defined looking back from the current micro instant. Thus, this macro state, while a function of several updates, can change every micro update, in a “sliding window” fashion.

To facilitate the distinction between a unit’s micro constituents *U*^*J*^ and its direct constituents *V* ^*J*^ —which may be meso units—we extend our definition of *J* above to be completely general, covering cases where *J* is a micro, meso, or macro unit. A unit *J* has five aspects:

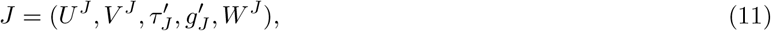

where *U*^*J*^ ⊆ *U* are its micro constituents, *V* ^*J*^ are its constituents (which may be micro or meso units) with current state *v*^*J*^ and state space 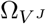, *W*^*J*^ is its background apportionment, which must contain the background apportionments of its constituents:

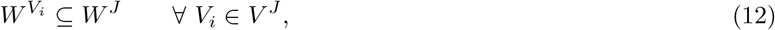

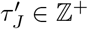 is the update grain over which *J*’s constituents are evaluated to define the state of *J*, and 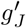 is a mapping from the states of *V* ^*J*^ over a sequence of 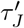 updates of *V* ^*J*^ to the state of *J* :

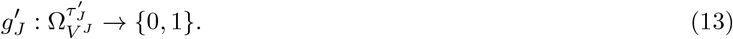

In general, there are 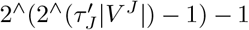 possible mappings from the state of constituents to the state of *J*. It is important to note that when *J* is constructed from a hierarchy of meso units of increasing grain, the update grain 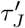 and the function 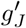 define a mapping across a *single* level of this hierarchy, from a sequence of states of *V* ^*J*^ to the state of *J*. If *V* ^*J*^ is a set of meso units, then 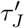 is the number of *meso* updates that define *J*’s state. There exist additional mappings between *V* ^*J*^’s constituents and *V* ^*J*^, and so on, down to the micro constituents *U*^*J*^ (with a corresponding nested sequence of background apportionments). Thus, in addition to the update grain of *J* in terms of its direct constituents 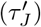, this hierarchical sequence of mappings can be used to define an update grain of *J* in terms of its *micro* constituents, which we label *τ*_*J*_. Similarly, we have a mapping *g*_*J*_ from sequences of micro states to the state of *J* :

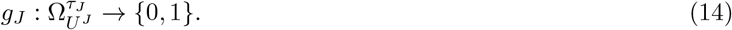

For example, in Figure 1B, 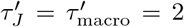 and *τ*_*J*_ = 8, because *J*’s state is defined over a sequence of 2 meso updates, each of which consists of 4 micro updates. Notice that unlike *τ* ^*′*^, *τ* is non-decreasing as a function of the level in the hierarchy.

#### 2.2.2 Applying the postulates to units

A complex, including its units, must comply with IIT’s postulates of physical existence [1]. Like complexes, macro units must have cause-effect power that is intrinsic, specific, irreducible, definite, and structured.

Consider the requirement for integration. Just like a complex, a candidate macro unit that does not satisfy integration, because it is reducible to causally independent subsets of micro units, can not truly exist as *one* unit. There is nothing unitary about it, except possibly from the extrinsic perspective of an experimenter. Pretending otherwise would be tantamount to building something (a macro unit, and then a complex) out of nothing (non-interacting micro units)(Fig. 2A). ^1^

**Figure 2:**
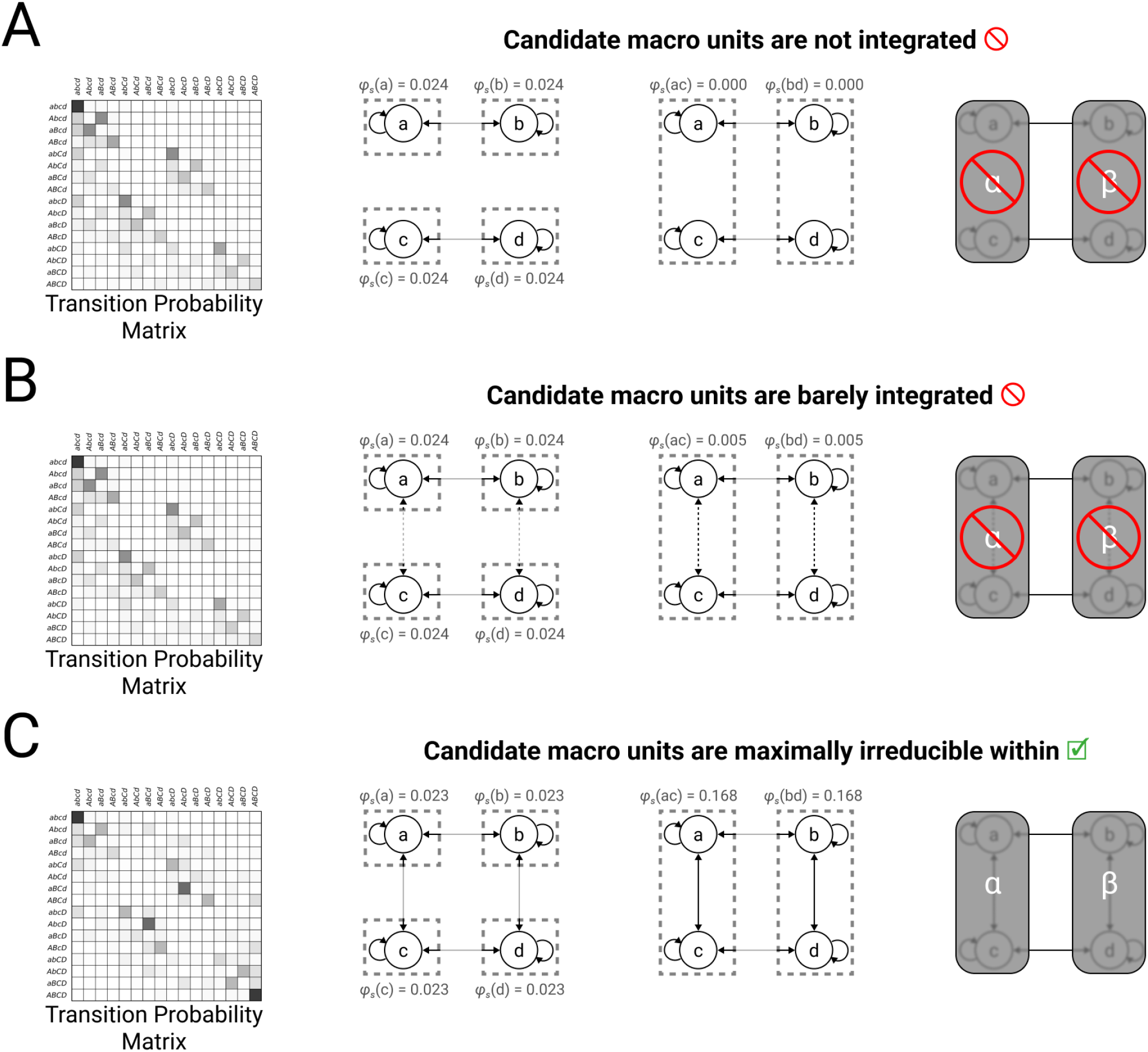
Out of nothing, nothing comes. Consider four micro units {*A, B, C, D*} in state (0, 0, 0, 0), constituting universe *U* with transition probability matrix *𝒯* _*U*_. For each micro unit *U*_*i*_, when all its inputs are 0, the probability that its state will be 1 after the next update is 0.05. This probability is increased by 0.05 if *u*_*i*_ itself is currently 1, and further increased by 0.6 if the state of *U*_*i*_’s horizontal neighbor is 1. (**A**) Consider the case where there are no connections between vertical neighbors (left). Each micro unit has *φ*_*s*_ = 0.024 on its own (middle left), while each pair of vertical neighbors has *φ*_*s*_ = 0 (middle right). Because each pair of vertical neighbors is reducible, they are not valid macro elements (right). (**B**) Consider the case where vanishingly weak connections are introduced between vertical neighbors, such that the probability that *u*_*i*_ will be 1 after the next update is increased by 0.01 if *U*_*i*_’s vertical neighbor is 1 (left). Although each pair of vertical neighbors is now very weakly integrated with *φ*_*s*_ = 0.005 (middle right), they are not maximally irreducible within (e.g., *φ*_*s*_(*a, b*) *< φ*_*s*_(*a*)). The conclusion is the same as for (A): the vertical neighbors are not valid macro elements (right). (**C**) Finally, consider the case where strong connections are introduced between vertical neighbors, such that the probability that *u*_*i*_ will be 1 after the next update is increased by 0.25 if *U*_*i*_’s vertical neighbor is 1 (left). Integration between vertical neighbors is now sufficiently strong (middle right) that the “maximally irreducible within” criterion is satisfied (middle right vs middle left), so we can consider macro elements built from vertical neighbors (right). There is no guarantee that the macro system consisting of these elements {*α, β*} is a complex, but at least we may evaluate that possibility (not shown).

Consider also the requirement for exclusion. Just as complexes must be definite, so must macro units: there must be a reason why a unit has the border it has. In the case of a complex, that reason is provided by the principle of maximal existence: the border is the one that yields maximal irreducibility. However, unlike complexes, intrinsic units only need to be maximally irreducible *within* (there cannot be any subset with higher integrated information) but not necessarily without (there can be supersets and/or parasets having higher integrated information). If intrinsic units were not required to be maximally irreducible within, one could treat as a macro unit a collection of nearly independent micro units, again building something out of “nearly nothing” (Figure 2). On the other hand, the units of a complex do not need to be maximally irreducible without, because the irreducibility to be maximized is that of the complex, rather than that of its units. Therefore, the borders of its intrinsic units should be those that maximize the complex’s *φ*_*s*_, rather than each unit’s *φ*. In summary, an intrinsic unit must be a maximally irreducible constituent of a complex (“maximally irreducible within”), rather than a complex itself (“maximally irreducible within and without”).

We now consider the requirements for intrinsic units more formally. To satisfy the intrinsicality, information and integration postulates, a unit *J* ∈ *S* with constituents *V* ^*J*^ (in current state *v*^*J*^) and background apportionment *W*^*J*^ must have cause-effect power that is intrinsic, specific, and irreducible:

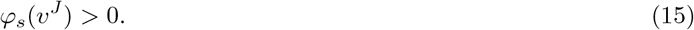

Moreover, to satisfy exclusion, *J* must have higher integrated information than any other valid system that could be constructed from its micro constituents *U*^*J*^ and background apportionment *W*^*J*^ :

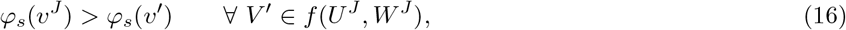

where *f* (*U*^*J*^, *W*^*J*^) is a set function that identifies all valid systems *V* ^*′*^ (ones that satisfy Eqs. 16 and 18) whose micro constituents are a subset of *U*^*J*^, and whose background apportionments are non-overlapping subsets of *W*^*J*^. The requirement that intrinsic units be maximally irreducible within applies whether macroing over units (Figure 3A-C), and/or over updates (Figure 3D), and applies to units at any grain.

Finally, consider a candidate system:

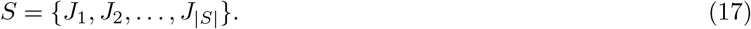

If the cause-effect power of a micro unit *U*_*i*_ ∈ *U* is not to be counted multiple times within *S*, (1) *U*_*i*_ cannot be a micro constituent of more than one macro unit; (2), it cannot be apportioned as background to more than one macro unit; and (3) it cannot be both a constituent of one macro unit and a mediator for another (this is analogous to the motivation for “screening-off” in other causal inference frameworks [24, 26, 10, 25]). This puts the following restriction on S:

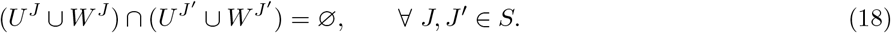

This restriction also ensures that the effects of multiple intrinsic units within a complex are integrated by other intrinsic units and not in the background. Because intrinsic units contribute cause-effect power to the complex but background conditions do not, (1) a complex can use another complex as background, but a unit cannot use another unit as background within a complex; (2) two complexes can use the same background, but two units cannot use the same background within the same complex.

**Figure 3:**
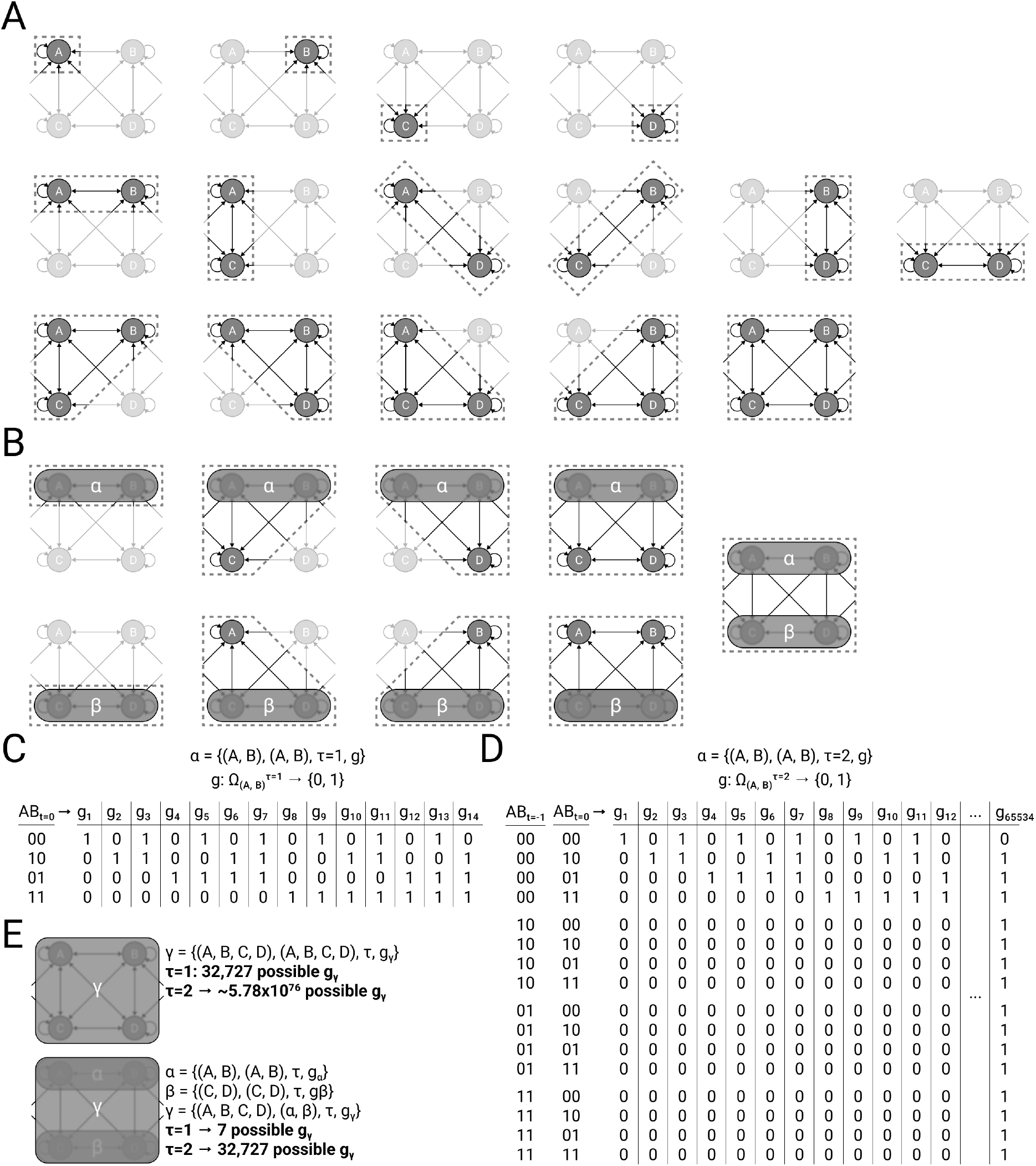
Defining macro units. Four micro units {*A, B, C, D*} are embedded within a larger universe *U*, with unspecified transition probability function 𝒯_*U*_. We wish to know if a macro unit *γ* can be built using micro constituents *U* ^*γ*^ = {*A, B, C, D*}, possibly with intermediate meso constituents *V* ^*γ*^ ≠ *U* ^*γ*^. (**A**) To ask if *γ* = {(*A, B, C, D*), …} is admissible as a macro unit, we must first check integrated information *φ*_*s*_ for every subset of micro units. (**B**) Suppose we find that {*A, B*} and {*C, D*} are maximally irreducible within (i.e., they satisfy Eqn. 16). This means that they are potential meso units, labeled *α* and *β* respectively. To continue verifying that *γ* = {(*A, B, C, D*), …} is admissible as a macro unit, we must now check integrated information for every subset of units that include *α* and *β* as well. (**C**) Let S be the macro system containing *α*. For a given candidate unit, say *α* = {(*A, B*), (*A, B*), *τ*_*α*_ = 1, *g*_*α*_)}, there are many potential mappings *g*_*α*_ from the states of *V* ^*α*^ = {*A, B*} over a sequence of *τ*_*α*_ = 1 updates to the state of *α*, but only one (here unspecified) will maximize *φ*_*s*_(*s*). (**D**) Same as (C), but over a sequence of *τ*_*α*_ = 2. Note that it is not only the ultimate state of the micro constituents that determine the macro unit’s state, but the precise sequence of micro states. (**E**) Depending on which of {*A, B, C, D*} or {*α, β*} (or some mixture) is maximally irreducible, *γ*’s constituents *V* ^*γ*^ might be {*A, B, C, D*} or {*α, β*} (or some mixture), which in turn will dictate the set of potential mappings from which *g*_*γ*_ can be defined, for any given *τ*. There are far fewer mappings that need to be considered for a macro unit whose constituents are meso units, because the mapping of the macro unit (*γ*) is constrained by the mappings of its meso constituents (*α, β*).

Having defined the criteria for an admissible system, the definition of a complex can be extended to arbitrary systems of units across grains. Let ℙ (*u*) be the set of all valid systems that can be defined from the universe *U* in state *u*. A system *S* = {*J*_1_, …, *J*_|*S*|_} in state *s* is a complex if it has more integrated information than any other admissible system that overlaps its micro constituents:

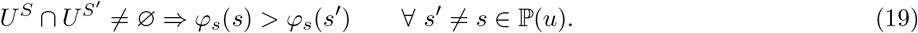

A consequence of this ‘maximally irreducible within’ requirement is that for a given set of micro constituents *U*^*J*^, whether an intrinsic unit *J* is built upon meso units (Figure 3E, bottom) or it is built directly “in one shot” upon the micro units (Figure 3E, top) depends on which definition of *V* ^*J*^ maximizes *φ*_*s*_(*v*^*J*^). In general, having finer (e.g., micro) constituents means having a larger number of mappings available to *J* with which to maximize *φ*_*s*_ at the macro grain, but makes it harder for *V* ^*J*^ to satisfy the requirement of being “maximally irreducible within.” For finer grain systems with a large number of constituents, having high *φ*_*s*_ requires that these units are both highly selective (to support intrinsic information), and have a connectivity structure without fault lines (to support integration) [19]. By contrast, coarser systems of units (defined from the same micro constituents) have fewer units, each of which can be flexibly defined through intermediate mappings to have the high selectivity and connectivity structure required to support high integrated information. Thus, although there is no strict requirement that a system of macro units be built up from meso units, there are good reasons to expect that many systems will have this property.

It is worth noting that whether a macro system is built up in levels, which precise macro and meso units it is built from, and which mappings define those units’ state, are all ultimately determined by what maximizes *φ*_*s*_ at each level of the hierarchy (the level that is maximally irreducible within). Thus, there is always a reason why a complex and its intrinsic units are precisely what they are: the principle of maximal existence.

Also note that the construction of *f* (*U*^*J*^, *W*^*J*^) is non-trivial, due to its dependence on *f* (*U*^*S*′^, *W*^*S*′^) for all *S*^*′*^ with *U*^*S*′^ ⊂ *U*^*J*^ ; the set of candidate systems depends on the set of admissible macro units, and the set of admissible macro units depends on the set of candidate systems within them (for satisfying “maximally irreducible within”). Practically, the sets need to be derived recursively. The starting point is that each micro unit *U*_*i*_ is a potential unit. The set of micro units then defines a set of candidate systems. Those candidate systems are then used as potential meso constituents for defining new potential units, which then leads to new candidate systems. The process can be repeated until convergence, which is guaranteed: the requirement of non-overlapping macro units ensures finite possible candidate systems.

#### 2.2.3 Assessing integrated information of macro systems: a conceptual overview

Having defined intrinsic units and the requirements for a system of intrinsic units to satisfy the postulates, we next outline a general framework for assessing *φ*_*s*_ that applies to any system, regardless of its units’ grains. In essence, we extend the definition of 𝒯_*c*_ and 𝒯_*e*_ to any system, whether it is constituted of micro units or macro units. These TPMs can then be used to compute *φ*_*s*_(*s*) as described in [1]. Here, we describe the process at a high level with some intuition for each step. Then, in Section 2.2.4, we will introduce some notation and provide a complete mathematical definition of the procedure.

For a system of macro units, we must define the intrinsic cause and effect TPMs (𝒯_*c*_ and 𝒯_*e*_) that describe their cause-effect power within the system, at their defined grains. Intuitively, one might consider using the universe’s micro TPM (𝒯_*U*_) to compute conditional probabilities between sequences of micro updates, and then *g*_*J*_ to map sequences of micro states to macro states. However, this process can expose micro cause-effect power in the macro TPMs (𝒯_*c*_ and 𝒯_*e*_) that is not intrinsic to the units at their defined grain (i.e., is extrinsic). To discount extrinsic cause-effect power, we employ a four step process (described in Section 2.2.4): (1) define modified transition probabilities between micro states; (2) use these to derive probabilities of sequences of micro updates; (3) causally marginalize background units; and (4) map the sequences of micro updates to macro states.

There are two situations in which system TPMs produced without the aforementioned modifications can lead to incorrect conclusions about the intrinsic, integrated cause-effect power of the system. The first is when the cause-effect power of a first macro unit over a third one is mediated by one or more micro constituents of a second macro unit. In this case, the cause-effect power of the these micro constituents will be counted twice: as belonging to both the first and second units. This is avoided by noising indirect pathways among macro units (see Eqn. 28 below).

The second situation is when a background unit outputs to two different macro units. In this case, the cause-effect power of the same micro unit would be counted twice, as a mediator for both macro units. This can be avoided by noising, for each background micro unit, the inputs from macro units other than the one it is apportioned to (see Eqn. 29 below).

After discounting the problematic interactions at the micro level, the modified transition probabilities can be used to compute the probability of sequences of micro states for the universe *U*, given a current micro state of the universe. This modified TPM only contains the cause-effect power at the micro grain that maps to cause-effect power at the macro grain. Once the transition probabilities have been extended to sequences, background units are causally marginalized conditional on the current state, resulting in conditionally independent transition probabilities between sequences of micro updates for the micro constituents of the substrate. Finally, the sequences of micro states are mapped to macro states to create 𝒯_*c*_ and 𝒯_*e*_.

The sequencing of these operations is important for the correct treatment of background conditions. First, causally marginalizing the background units happens after the transition probabilities are extended to sequences of micro updates. The conditional causal marginalization ensures that the analysis starts from the current state of background units (the context for the system’s cause-effect power), but does not keep them fixed throughout the analysis. This allows the background conditions to ‘percolate’ and mediate interactions among macro units, rather than being absolutely frozen in the current state. Second, the causal conditioning of background units should occur before sequences of micro states are mapped into macro states. This is because the background units must be treated at the micro grain; they are extrinsic to the system, and do not exist as macro units from the intrinsic perspective of the system (even if they are macro units within another complex).

#### 2.2.4 Assessing integrated information of macro systems: mathematical framework

When dealing with macro update grains, we require additional notation. Let _*t*_*u* ∈ Ω_*U*_ be the state of *U* at update *t*. Similarly, let _*t*_*u*_*i*_ be the state of micro unit *U*_*i*_ ∈ *U* at update *t*. Let *t* = 0 denote the current micro update, so the current micro state of the universe is _0_*u*. Negative prescripts (*t <* 0) index the updates that led to the current micro state, and positive prescripts (*t >* 0) index the updates that follow from the current micro state. Finally, let _*t*_*ů* be a sequence of *τ* micro states leading to _*t*_*u*:

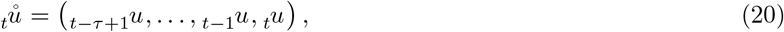

and let _*t*_*ũ* be a sequence of *τ* micro states following _*t*_*u*:

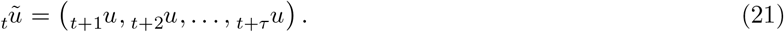

For example, _0_*ů* is the sequence of *τ* micro states leading to the current micro state _0_*u*. To reduce notational burden, we omit the prescripts and use *ů* and *ũ* when referring to generic *τ* -length sequences that lead to or follow from a generic *u*.

For a macro unit *J* with macro update grain *τ*_*J*_, its current macro state *j* depends on the previous *τ*_*J*_ states of its micro constituents *U*^*J*^ (Fig. 1B):

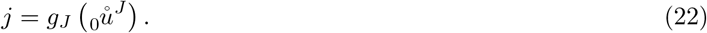

Consider a system *S* = {*J*_1_, …, *J*_|*S*|_} in state *s* = (*j*_1_, *j*_2_, …, *j*_|*S*|_). Each unit *J*_*i*_ (whether micro or macro) has micro constituents 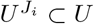, constituents 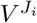, background apportionment 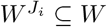, an update grain 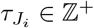, and a mapping 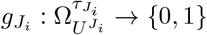. We denote the system’s micro constituents as:

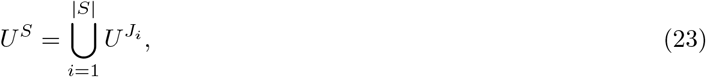

and its background units as:

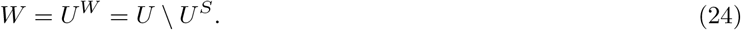

*W* and *U*^*W*^ are equivalent (unlike *U*^*S*^ and *S*, for example) and may be used interchangeably where notationally convenient, because background units are always treated at the micro grain. The background apportionments for the system, *W*^*S*^ ⊆ *W*, are simply the (non-overlapping, see Eqn. 18) apportionments of its constituents:

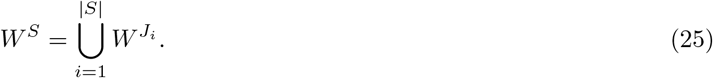

After discounting connections extrinsic to the system (see below), the micro units in 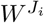 will receive intact inputs from 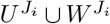, but not from other units. A consequence is that the background units cannot integrate cause-effect power.

Next, we define generalized cause and effect TPMs *𝒯*_*c*_ and *𝒯*_*e*_ for *S*, from which the rest of the framework can be applied as usual. The process proceeds in four steps: (1) starting from the universe TPM *𝒯*_*U*_, for each macro unit *J*, discount any connections that are extrinsic to the system (e.g., cause-effect power from micro grains that does not map to macro grains), yielding modified transition probabilities 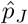 between micro states; (2) extrapolate the modified transition probabilities 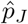 into probabilities for sequences of micro updates given a current micro state; (3) causally marginalize the background (*W*) conditional on the current sequence of micro states; and (4) use the mappings 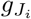 to compress the micro state-by-sequence transition probabilities for each macro unit into its macro state-by-state transition probabilities, finally combining these macro transition probabilities for each unit to get macro system TPMs 𝒯_*c*_ and 𝒯_*e*_. When no macroing is performed (i.e., all 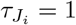 and all 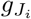 are identity functions), (1), (2), and (4) are trivial, and 𝒯 _*c*_ and 𝒯 _*e*_ work out to be (micro) TPMs as defined in [1].

**Step 1: Discounting connections extrinsic to the system**. We wish to define modified state transition probabilities that reflect the effects of discounting certain micro connections. The specific micro connections to be discounted will depend on the macro unit being updated. For example, if we are updating the state of *J*_1_ then connections from *J*_2_ to *J*_1_ are left intact, but if we are updating the state of *J*_3_, those connections are noised to prevent *J*_1_’s micro constituents from mediating other units’ effects. To have connections selectively discounted, we will define a different 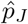 for each macro unit *J* ∈ *S* to be updated, and then combine them later with a product. The product removes any cause-effect power from the TPM whose source is correlations among units due to common input from background units or micro constituents, leaving intact the direct cause-effect power among units.

For a given *J* ∈ *S* and *U*_*i*_ ∈ *U*, we would like to define:

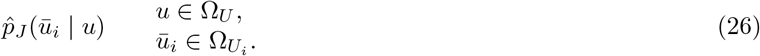

If *U*_*i*_ ∈ *U*^*J*^ (*U*_*i*_ is a constituent of the to-be-updated macro unit), then no connections are discounted:

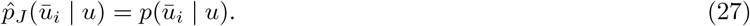

If *U*_*i*_ ∈ *U*^*S*^ *\U*^*J*^ (in the system, but not a constituent of *J*) or *U*_*i*_ ∈ *W \W*^*S*^ (a background unit that is not apportioned to any system unit), then all connections should be discounted:

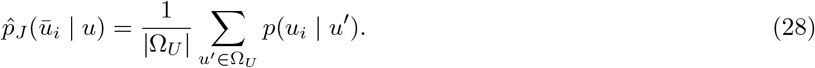

This ensures that the micro units that constitute macro units do not have a second role as mediators of other macro units’ effects (e.g., *J*_1_ effects *J*_3_ through *J*_2_’s micro constituents).

If 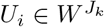, then all connections from 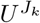 and 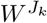 should be kept intact, but all other connections should be noised (allowing 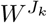 to mediate *J*_*k*_’s effects, but no other units’ effects):

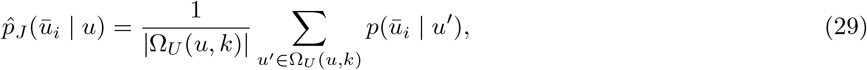

where 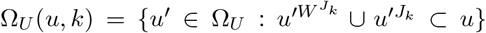 is the set of all universe states where the state of *J*_k_’s micro constituents 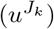 and background apportionment 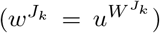 are consistent with *u*. Averaging over system states discounts (noises) all micro connections to 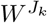 from outside 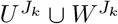.

The modified unit probabilities, 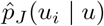, can then be combined to create a modified universe TPM that contains only the connections required to update the state of *J*,

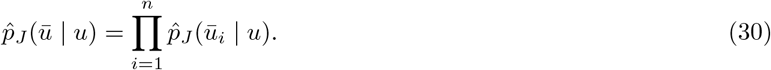

**Step 2: Obtaining probabilities for sequences of micro updates**. The modified transition probabilities can now be used to compute the probability of any sequence of *τ*_*J*_ micro states _*t*_*ũ*, conditional on any given state _*t*_*u*:

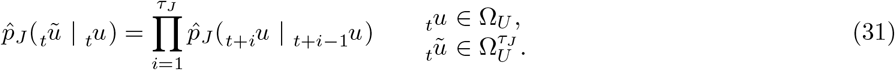

Note that 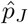 may denote state-to-state transition probabilities (as in Eqn. 30, or the right side of Eqn. 31) or state-to-sequence transition probabilities (as in the left side of Eqn. 31), depending on its arguments (e.g., 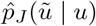 vs. 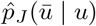).

**Step 3: Causally marginalizing the background**. For each unit’s modified transition probability function 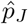, cause and effect versions 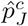 and 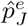 are computed by causally marginalizing the background units conditional on _0_*ů*.

This entails taking a weighted average of state-to-sequence transition probabilities over potential background states, weighting each by the probability *q*_*c/e*_(*w* | _0_*ů*) of that background state given the previous *τ*_*J*_ micro states of the universe:

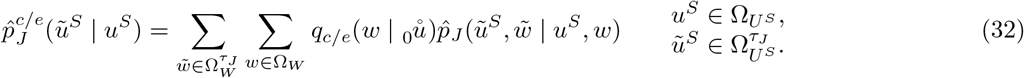

To get the weights *q*_*c/e*_, suppose _0_*ů* is the sequence of micro states corresponding to the current macro state. For evaluating effects, the current background state _0_*w* gets full weight (there is no uncertainty about the current micro state given _0_*ů*). We directly condition on the current micro state to get:

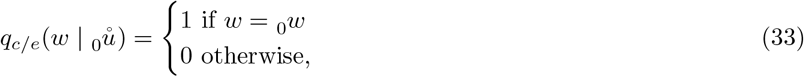

For evaluating causes, 𝒯_*U*_ is used in combination with Bayes’ rule to determine a probability distribution for 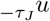, conditional on the current macro state. Because of the Markov property, only the distribution for 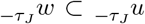 is required to weight 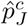, and only the earliest micro state in the sequence (i.e., only 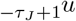) is required for the Bayesian computation. A uniform marginal distribution of the previous updates is assumed (i.e., maximum uncertainty about prior states, see also Section 2.1):

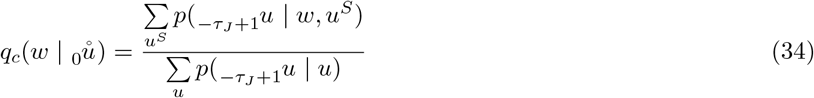

**Step 4: Compressing state-by-sequence transition probabilities into macro-state TPMs**. Finally, micro state-to-sequence transition probabilities are mapped to probabilities of individual macro updates. First, for each macro unit, we obtain the probability of transitioning to each of its macro states, given each possible micro state of the system:

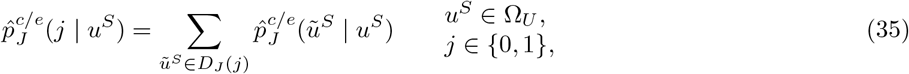

where *D*_*J*_ (*j*) is the set of sequences of micro states that are mapped to *J* = *j*:

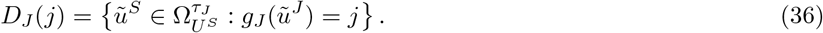

Then we map each current micro system state *u*^*S*^ to corresponding macro states. Generally, each current micro state will be mapped to different current macro states some proportion of the time, depending on the sequence of micro states that led to it, and on the mapping *g*_*J*_. For each macro system state *s*:

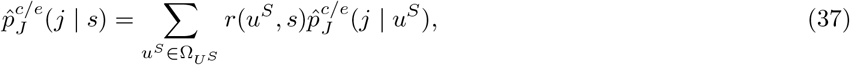

where *r*(*u*^*S*^, *s*) is the proportion of sequences of micro states _*t*_*ů*^*S*^ that end with *u*^*S*^ (i.e., 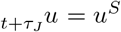), among all the sequences of micro states that get mapped to system state *s*:

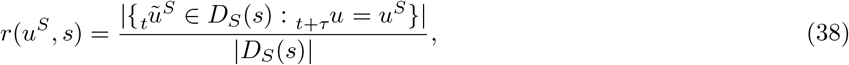

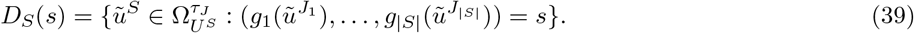

Operationally, this amounts to a procedure where perturbing a macro unit into its state is achieved by perturbing its micro constituents into all possible micro state sequences that map to the corresponding macro state, with equal probability.

The probability functions 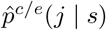 define the probability of the future macro state of *J* given the current macro state of *S*. Finally, we combine the functions for each *J* as a product (establishing conditional independence and removing any correlations due to extrinsic factors):

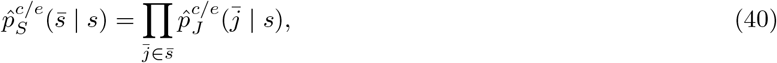

where 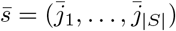.

For a system of macro units, its *φ*_*s*_ value is computed from the cause and effect TPMs of *S*:

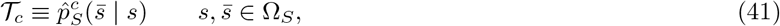

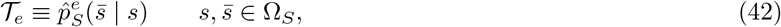

as described in [1]. It is worth noting that once 𝒯_*c*_ and 𝒯_*e*_ have been derived, there is no further reference to the background units, the grain of the units, or their micro constituents. For example, to assess the intrinsic information of the system, the macro units are perturbed, equally likely, into all possible states, regardless of whether or not this corresponds to a uniform distribution for the states of the micro constituents. The TPMs are taken to describe the intrinsic cause-effect power of the system’s units, at their particular grain.

## 3 Results

In this section, the framework is applied to two example systems. The examples demonstrate that intrinsic cause-effect power can be higher for a system of macro units than for any system of the corresponding micro units, extending results from earlier work [15, 18] to the updated framework [1]. Computations of integrated information were performed using PyPhi [21]. In what follows, we omit the state as input to *φ*_*s*_ (e.g., *φ*_*s*_({*A, B*})) when the state of the units can be inferred from the context of the example.

### 3.1 Example 1

Consider four micro units {*A, B, C, D*} in state (0, 0, 0, 0), constituting universe *U* with transition probability matrix 𝒯_*U*_ (Figure 4A). Each micro unit *U*_*i*_ has the same function: When all its inputs are 0, the probability that its state will be 1 after the next update is 0.05. This probability is increased by 0.01 if *u*_*i*_ itself is currently 1. Thus, there is a very weak tendency for a unit that is 1 to remain 1. The probability that *u*_*i*_ will be 1 after the next update is increased by 0.1 if the state of *U*_*i*_’s horizontal neighbor is 1. For example, *A* is more likely to be 1 after the next update if *B* is currently 1. Finally, the probability that *u*_*i*_ will be 1 after the next update is increased by 0.8 if *both* its vertical neighbor and its diagonal neighbor are currently 1. For example, *A* is very likely to be 1 after the next update if both *C* and *D* are currently 1. Thus, each micro unit approximates a noisy logical AND function over its vertical and diagonal neighbors, with a weak independent influence from its horizontal neighbor, and very weak self-influence. Because each unit’s future state is mostly dictated by its vertical and diagonal neighbors (e.g., *A*’s future state depends most heavily on the current states of *C* and *D*), and because horizontal neighbors share the same vertical and diagonal neighbors, (e.g., both *A* and *B* are dominated by *C* and *D*), we expect that macroing horizontal neighbors into macro units will reduce the indeterminism and degeneracy associated with the micro units (by combining multiple low probability micro states into a single higher probability macro state), and thereby increase cause-effect power [16, 15, 18].

**Figure 4:**
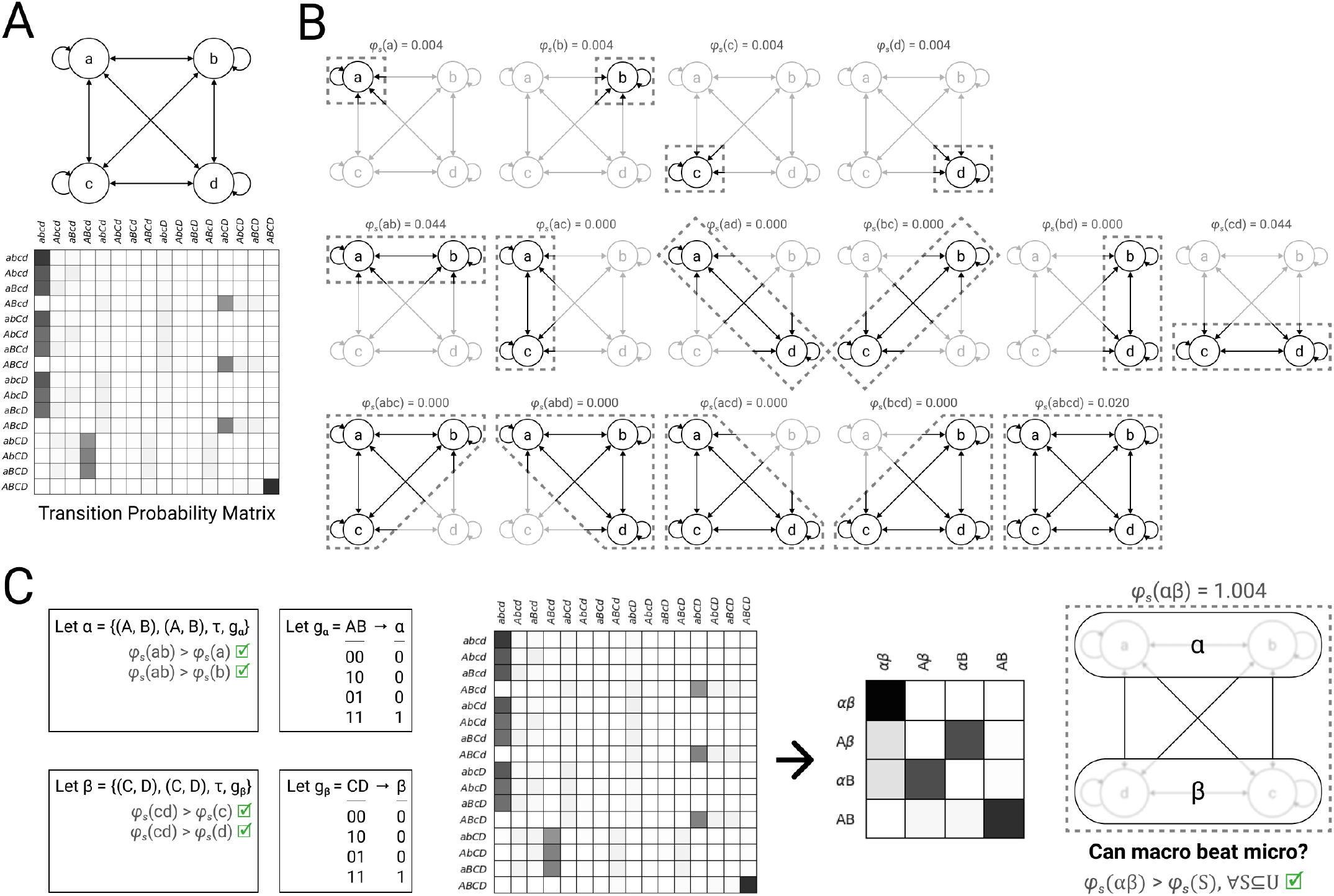
Example 1. (**A**) Consider four micro units {*A, B, C, D*} in state (0, 0, 0, 0), with transition probability matrix 𝒯_*U*_. For illustrative purposes, capitalization denotes the state of each unit, both in causal network diagrams and transition probability matrix state labels (e.g., state (0, 1, 0, 0) is written *aBcd*). (**B**) System integrated information *φ*_*s*_(*s*) must be checked for each subset of micro units *S* ∈ ℙ ({*A, B, C, D*}). Greyed-out units are background. Notice that {*A, B*} and {*C, D*} are maximally irreducible within. In the case of {*A, B*}: *φ*_*s*_({*A, B*}) = 0.044, greater than either *φ*_*s*_(*A*) = 0.004 or *φ*_*s*_({*B*}) = 0.004. (**C**) Since {*A, B*} and {*C, D*} are maximally irreducible within, we may consider their potential macro units, labeled *α* and *β* respectively. One possible pair of mappings for these macro units are *g*_*α*_ and *g*_*β*_, resulting in macro TPM 𝒯^*S*^. This candidate system {*α, β*} in state (0, 0) (given by *g*_*α*_, *g*_*β*_) has system integrated information *φ*_*s*_ = 1.004, greater than any of the micro level candidate systems in (B). Thus, although we would have to check all other valid macro unit definitions and mappings in order to determine whether this macro system is *maximally* irreducible relative to all others, we can conclude that intrinsic cause-effect power will be higher at a macro level than at the micro level—we know that we can do at least as well as *φ*_*s*_ = 1.004.

To confirm this intuition, we first assess the system integrated information *φ*_*s*_ of all possible candidate systems of micro units (Figure 4B). At this micro level, the two candidate systems with the most irreducible cause-effect power are {*A, B*} and {*C, D*}, both with *φ*_*s*_ = 0.044. Because these candidate systems are maximally irreducible within (i.e., *φ*_*s*_({*A, B*}) *> φ*_*s*_(*s*) *∀S* ⊆ {*A, B*}), they satisfy Eqn. (16) and can be considered as macro units. Notice that although {*A, B, C, D*} as a whole has irreducible cause-effect power (*φ*_*s*_ = 0.020), it is not maximally irreducible within (e.g., *φ*_*s*_({*A, B*}) *> φ*_*s*_({*A, B, C, D*})) and *cannot* be considered as a macro unit.

Let macro unit *α* be defined from micro constituents {*A, B*}, and *β* from {*C, D*}. There are 14 possible mappings for each macro unit (Figure 3C). In particular, the mapping shown in Figure 4C seems promising, because it ought to decrease both the indeterminism and the degeneracy that are present in the micro system. This class of mapping, in which the state of the macro unit is a simple function of the number of constituents in state 1, has also been referred to as “coarse-graining” [16, 15]. Coarse-graining corresponds to the typical notion of a macro state in statistical physics [18]. Under the mapping shown in Figure 4C, each macro unit’s state is 1 if-and-only-if both its micro constituents are 1. When one macro unit is 1, odds are that the other macro unit will be 1 after the next update. When one macro unit is 0, odds are that the other macro unit will be 0 after the next update. Thus, the macro system behaves something like two reciprocally connected COPY gates, with some additional complexity provided by the connections between horizontal neighbors at the micro level. This is reflected in the macro system’s TPM (Figure 4C, middle). Indeed, when we measure the system integrated information of {*α, β*}, we find *φ*_*s*_({*α, β* }) = 1.004, demonstrating that this system of macro units has more irreducible, intrinsic cause-effect power than any candidate system built without macro units (Figure 4C, right).

#### 3.2 Example 2

Consider eight micro units {*A, B, C, D, E, F, G, H*} in state (1, 1, 1, 1, 1, 1, 1, 1), constituting universe *U* with transition probability matrix 𝒯_*U*_ (Figure 5A). The left half of the system and the right half of the system ({*A, B, C, D*} and {*E, F, G, H*}, respectively) are mirror images of each other, so for simplicity consider the left half. For every unit *U*_*i*_, the probability that its state will be 1 after the next update is marginally higher if its current state is 1. *C* approximates a noisy logical OR function of *A* and *B*, which in turn approximate a noisy logical COPY function of *C*’s image *G*. When *A, B*, or *C* are 1, the probability that *D* is 1 after the next update increases linearly. *D*’s current state also has weak influence on the future state of *A* and *B*. Roughly speaking, we can think of the two halves of the system as copying each other’s state, but whereas a disruption to any of the connections *within* either half will moderately disrupt this function, a disruption to any of the connections *between* halves will severely disrupt it. It is reasonable to expect that macroing the left half of the system and the right half of the system into separate macro units, and treating the macro units’ states as a simple function of *C* and *G*’s states, will increase intrinsic cause-effect power. This class of mapping, in which the state of the macro unit is determined only by the state of specific constituents, ignoring others, has also been referred to as “black-boxing.” Black boxes correspond to the typical notion of macro units in the special sciences, because they are constituted of heterogeneous micro units that are often compartmentalized and have highly specific functions, which would be muddled by averaging [18].

**Figure 5:**
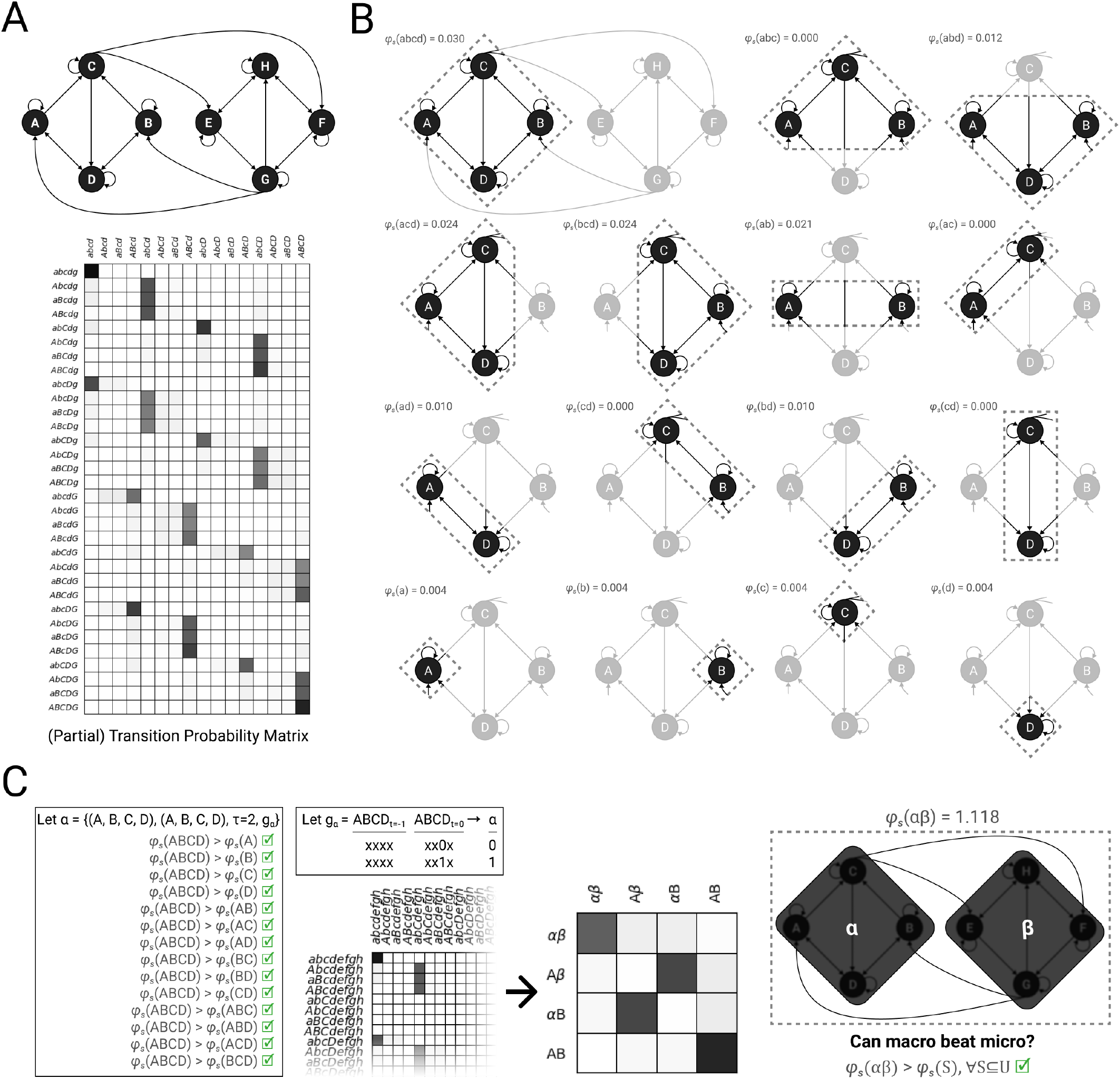
Example 2. (**A**) Consider eight micro units {*A, B, C, D, E, F, G, H*} in state (1, 1, 1, 1, 1, 1, 1, 1), with transition probability matrix (TPM) 𝒯_*U*_. Because of space limitations in this and subsequent panels, we illustrate some analysis steps for the left half of the system only (i.e., {*A, B, C, D*}), but all calculations were done using the full eight-unit system. For example, although the full TPM is used for all calculations, a partial TPM illustrating the behavior of {*A, B, C, D*} is shown here. Rows are past system states and columns are future states. (**B**) System integrated information *φ*_*s*_(*s*) must be checked for each subset of micro units *S* ∈ P({*A, B, C, D, E, F, G, H*}). Here, because of space limitations, we illustrate checks for *S* ∈ ℙ ({*A, B, C, D*}). Notice that {*A, B, C, D*} is maximally irreducible within. (**C**) Since {*A, B, C, D*} is maximally irreducible within, we may consider its potential macro unit, labeled *α*. One possible mapping for *α* with *τ* = 2 is shown. Since the full system is symmetric, {*E, F, G, H*} can be considered as a potential macro unit *β* with analogous *g*_*β*_, resulting in macro TPM 𝒯^*S*^. This candidate system {*α, β*} in in state (1, 1) (given by *g*_*α*_, *g*_*β*_) has system integrated information *φ*_*s*_ = 1.118, greater than all of the micro level candidate systems (max *φ*_*s*_ = 0.135, not shown, but see (B) for a subset). Thus, although we would have to check all other valid macro unit definitions and mappings in order to determine whether this macro system is *maximally* irreducible relative to all others, we can conclude that intrinsic cause-effect power will be higher at a macro level than at the micro level—we know that we can do at least as well as *φ*_*s*_ = 1.118.

To compare the cause-effect power of the micro and macro systems, we first assess the system integrated information *φ*_*s*_ of all possible candidate systems of micro units (Figure 5B). Note that, in addition to the candidate systems shown in Figure 5B, all candidate systems of five units (e.g., {*A, B, C, D, E*}), six units (e.g., {*A, B, C, D, E, F* }), and seven units (e.g., {*A, B, C, D, E, F, G*}) were evaluated (not shown). At the micro grain, the maximum value of *φ*_*s*_ is 0.135 (*S* = {*A, C, E, G*} and symmetric systems). At the micro grain, the two candidate systems that we hypothesized would make good macro units ({*A, B, C, D*} and {*E, F, G, H*}) are maximally irreducible within, with *φ*_*s*_ = 0.030. Because these candidate systems are maximally irreducible within (i.e., *φ*_*s*_({*A, B, C, D*}) *> φ*_*s*_(*s*) ∀*s ⊂* {*A, B, C, D*}), they satisfy condition (16) and can be considered as macro units.

Let macro unit *α* be defined from micro constituents {*A, B, C, D*}, and *β* from {*E, F, G, H* }. Our mapping of interest, where the state of *α* is dictated by the state of its output unit *C* over two micro updates (*τ* = 2), is shown in Figure 5C. Under this mapping, the macro system behaves something like two reciprocally connected COPY gates, with some additional complexity provided by the connections within each macro unit. This is reflected in the macro system’s TPM (Figure 5C, middle), which is very similar to the macro TPM obtained in the previous example (Figure 4C, middle). Indeed, when we measure the system integrated information of {*α, β*}, we find *φ*_*s*_({*α, β*}) = 1.118, demonstrating that this system of macro units has more irreducible, intrinsic cause-effect power than any candidate system built without macro units (Figure 5C, right).

## 4 Discussion

In this work, we extend IIT’s mathematical framework for assessing cause-effect power (IIT 4.0 [1]) to systems of macro units, explaining how this framework formalizes IIT’s postulates of existence, intrinsicality, information, integration, exclusion, and composition. Based on this framework, we further demonstrate that macro grain systems (systems constituted of one or more macro units) can have higher cause-effect power than the corresponding micro grain systems, as measured by system integrated information (*φ*_*s*_).

IIT’s existence postulate requires that macro units have cause-effect power (that they “take and make a difference”), as established operationally by manipulating and observing their state. The intrinsicality, information, integration, and exclusion postulates require that the cause-effect power of macro units be intrinsic, specific, irreducible (*φ*_*s*_ *>* 0), and definite in grain. Based on IIT’s principle of maximal existence (among competing existents, the one that actually exists is the one that exists the most), the grain is such that (i) each unit is maximally irreducible “within” (it has greater *φ*_*s*_ than any combination of its constituents) and (ii) the units taken together maximize irreducibility of *φ*_*s*_ over their substrate. These then constitute the complex’ “intrinsic units.” From the perspective of the complex whose structure they compose, intrinsic units have no internal structure of their own, and exist in one of two alternative macro states.

Searching across grains for maxima of *φ*_*s*_ assumes that cause-effect power can be highest at macro grains [16, 15, 18]. The examples presented in this work employing the IIT 4.0 framework demonstrate that this is indeed possible. Specifically, we show that a macro system can have greater cause-effect power than the corresponding micro system if it is associated with reduced indeterminism and degeneracy of state transitions, such that the selectivity of causes and effects is correspondingly increased (see also [16, 15, 12]). In IIT 4.0, this increased selectivity is captured naturally by *φ*_*s*_ because of its formulation in terms of intrinsic information [8, 7, 19]. Increased selectivity can arise at macro grains because the analysis of cause-effect power treats each macro state of the macro units as equally likely, corresponding to a non-uniform distribution of micro states. Moreover, a system of macro units can have greater cause-effect power than the corresponding micro units if integration is higher at the macro level (see also [18]).

To achieve a high value of *φ*_*s*_, systems of any grain must balance integration with differentiation. Whether *φ*_*s*_ will increase with a larger number of units depends on a balance between how much additional cause and effect information the system can specify (because its state repertoire has expanded), how much the selectivity of causes and effects within the system is reduced (because cause and effect information is spread over additional states, even more so if the additional units bring increased noise), and how well integrated the additional units are with the rest of the system [8, 7, 19]. Thus, a system of many units can only “hang together well” as an intrinsic entity if its units are themselves highly integrated and are appropriately interconnected, say as a dense, directed lattice [1]. We conjecture that macro units built upon a hierarchy of meso units may play a crucial role in allowing large systems to exist as maxima of intrinsic, irreducible cause-effect power. Hierarchies of this sort appear to be a common feature of biological systems.

In general, macro grains with *φ*_*s*_ values higher than *most* finer or coarser grains—that is, local or “extrinsic” maxima of integration and causal efficacy [16]—are likely to capture relevant levels of substrate organization by “carving nature at its joints.” In the brain, for example, these might correspond to proteins, ion channels, organelles, synaptic vesicles, synapses, neurons, groups of tightly interconnected neurons, and so on. Such “extrinsic units,” well-suited to manipulations and observations by neuroscientists, are critical for understanding how the system works. However, according to IIT, there is a critical difference between these locally maximal grains and the absolute maximal grain whose “intrinsic units” maximize *φ*_*s*_ within and without: only the latter constitutes the substrate of consciousness and contributes to the way the experience feels—all other levels of organization do not exist from the intrinsic perspective.

Another consequence of IIT’s intrinsic framework has to do with update grains. Assuming the grain of intrinsic units is that of neurons, the update grain might be, for instance, on the order of 30 milliseconds (in line with estimates of the duration below which non-simultaneous sensory stimuli are perceived as being simultaneous, or changing stimuli are perceived as static, [30]). From the extrinsic perspective of an experimenter, several update grains may be critical to understand different kinds of causal interactions—finer grains for events such as ion channel opening, quantal release of transmitters, and the like—and longer grains for low-frequency synchronization, the induction of plastic changes, and so on. But again, while these faster and slower time scales are critical for understanding how the system works, only one time scale matters intrinsically—from the perspective of the conscious subject. Accordingly, IIT predicts that experience should only change if there is a change in the state of intrinsic units at their intrinsic update grain. Any other changes will affect the brain, but not experience. Even more stringently, the requirement that intrinsic units have binary macro states implies that any change in their micro state that does not translate into a switch of their macro state will not affect experience. For example, changes in the timing of neuronal firing, or in the rate of firing, may have clear-cut effects on the rest of the brain, but if they map onto the same intrinsic macro state, they will not have effects on the experience.

For IIT, physical existence is cause-effect power, with no further need for intrinsic properties. IIT’s analysis of cause-effect power starts from a causal model of a micro-physical substrate, which is defined by its transition probabilities, 𝒯_*U*_ (Eqn. 1). The substrate TPM is taken to be a complete description of the substrate. The examples presented were analyzed based on the assumption that the substrate TPM was fully known and strictly stationary, with the goal of demonstrating the self-consistency of IIT’s approach and highlighting some of its consequences. In practice, the determination of complexes and their intrinsic units cannot be based on a full knowledge of the micro-physical TPM. At most, the theoretical principles outlined here can serve as a heuristic guidance for pointing to candidate complexes and intrinsic units, at the expense of many assumptions and approximations. Stationarity of the TPM over macro states is also merely a convenient assumption. In general,𝒯 𝒯_*U*_ is expected to evolve at every update, in accordance with IITs *principle of becoming* (“powers become what powers do,” to be considered in future work [28]).

## Data Availability

No new data were generated or analysed in support of this research. All code used to obtain figures and examples can be found at [13].

## Funding

This project was made possible through the support of a grant from Templeton World Charity Foundation (TWCF0216, G.T.). In addition, this research was supported by the David P White Chair in Sleep Medicine at the University of Wisconsin-Madison, by the Tiny Blue Dot Foundation (UW 133AAG3451; G.T.), and by the Natural Sciences and Engineering Research Council of Canada (NSERC; RGPIN-2019-05418; W.M.). The funders had no role in study design, data collection and analysis, decision to publish, or preparation of the manuscript.

## Competing interests

G.T. holds an executive position and has a financial interest in Intrinsic Powers, Inc., a company whose purpose is to develop a device that can be used in the clinic to assess the presence and absence of consciousness in patients. This does not pose any conflict of interest with regard to the work undertaken for this publication.

## Author contributions

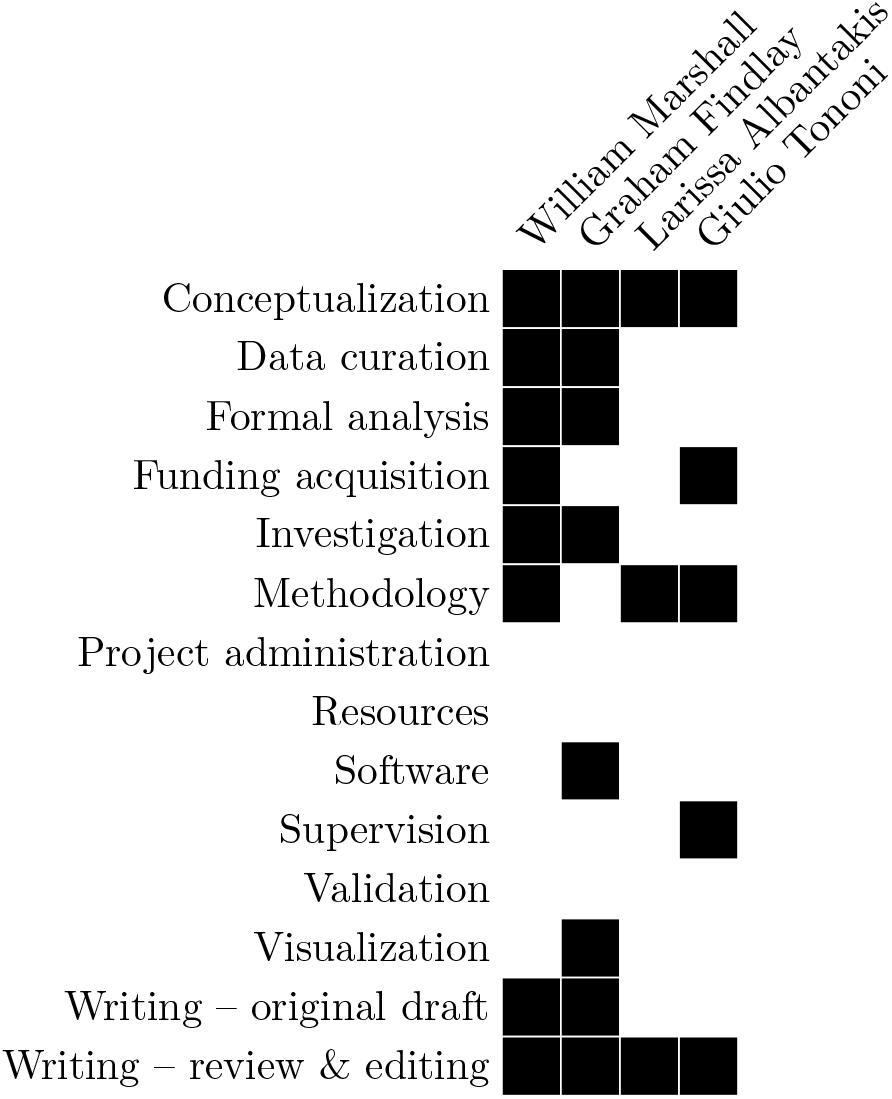

## Footnotes

1 As mentioned above, previous versions of the theory [18] failed to impose this requirement on macro units directly—only on the system—which required partitions testing system irreducibility to be performed at the micro level, often cutting through macro units. Although this did limit the gerrymandering of macro units and complexes illustrated in Fig. 2, it contradicted the premise that from the intrinsic perspective of the system, it exists as a collection of macro (rather than micro) units.

## Notes

### Summary of Updates

Title changed; figures revised; extended results.

## References

[1] Larissa Albantakis, Leonardo Barbosa, Graham Findlay, Matteo Grasso, Andrew M. Haun, William Marshall, William GP. Mayner, Alireza Zaeemzadeh, Melanie Boly, Bjorn E. Juel, Shuntaro Sasai, Keiko Fujii, Isaac David, Jeremiah Hendren, Jonathan P. Lang, and Giulio Tononi. Integrated information theory (iit) 4.0: formulating the properties of phenomenal existence in physical terms. PLoS Comp. Biol., 2023.

[2] Larissa Albantakis, Arend Hintze, Christof Koch, Christoph Adami, and Giulio Tononi. Evolution of Integrated Causal Structures in Animats Exposed to Environments of Increasing Complexity. PLoS computational biology, 10(12):e1003966, ec 2014.

[3] Larissa Albantakis, William Marshall, Erik Hoel, and Giulio Tononi. What caused what? A quantitative account of actual causation using dynamical causal networks. Entropy, 21(5):459, may 2019.

[4] Larissa Albantakis, Robert Prentner, and Ian Durham. Measuring the integrated information of a quantum mechanism. Entropy, 25, 2023.

[5] Nihat Ay and Daniel Polani. Information Flows in Causal Networks. Advances in Complex Systems, 11(01):17– 41, feb 2008.

[6] David Balduzzi and Giulio Tononi. Integrated information in discrete dynamical systems: motivation and theoretical framework. PLoS Comput Biol, 4(6):e1000091, jun 2008.

[7] Leonardo S Barbosa, William Marshall, Larissa Albantakis, and Giulio Tononi. Mechanism Integrated Information. Entropy, 23(3):362, March 2021.

[8] Leonardo S Barbosa, William Marshall, Sabrina Streipert, Larissa Albantakis, and Giulio Tononi. A measure for intrinsic information. Scientific Reports, 10(1):18803, 2020.

[9] Melanie Boly, Marcello Massimini, Naotsugu Tsuchiya, Bradley R Postle, Christof Koch, and Giulio Tononi. Are the neural correlates of consciousness in the front or in the back of the cerebral cortex? clinical and neuroimaging evidence. Journal of Neuroscience, 37(40):9603–9613, 2017.

[10] Robert Brandon. The Levels of Selection. PSA: Proceedings of the Biennial Meeting of the Philosophy of Science Association, 1982:315–323, 1982.

[11] Robert Chis-Ciure, Jeremiah Hendren, Matteo Grasso, Bjørn Erik Juel, and Giulio Tononi. FAQ: If IIT assumes ’physicalism,’ does this make it a materialist theory of consciousness? http://www.iit.wiki/faqs/philosophy, June 2024.

[12] Renzo Comolatti and Erik Hoel. Causal emergence is widespread across measures of causation, February 2022.

[13] Graham Findlay and William Marshall. https://github.com/CSC-UW/Marshall_et_al_2024/releases/tag/ v0.1.0. doi:10.5281/zenodo.11211436, 2024.

[14] Andrew M Haun and Giulio Tononi. Why Does Space Feel the Way it Does? Towards a Principled Account of Spatial Experience. Entropy, 21(12):1160, nov 2019.

[15] Erik P. Hoel, Larissa Albantakis, William Marshall, and Giulio Tononi. Can the macro beat the micro? Integrated information across spatiotemporal scales. Neuroscience of Consciousness, 2016(1), 2016.

[16] Erik P. Hoel, Larissa Albantakis, and Giulio Tononi. Quantifying causal emergence shows that macro can beat micro. PNAS, 110(49):19790–19795, nov 2013.

[17] Dominik Janzing, David Balduzzi, Moritz Grosse-Wentrup, and Bernhard Schölkopf. Quantifying causal influences. The Annals of Statistics, 41(5):2324–2358, oct 2013.

[18] William Marshall, Larissa Albantakis, and Giulio Tononi. Black-boxing and cause-effect power. PLOS Computational Biology, 14(4):e1006114, apr 2018.

[19] William Marshall, Matteo Grasso, William GP Mayner, Alireza Zaeemzadeh, Leonardo S Barbosa, Erick Chastain, Graham Findlay, Shuntaro Sasai, Larissa Albantakis, and Giulio Tononi. System Integrated Information. Entropy, 25, 2023.

[20] William Marshall, Hyunju Kim, Sara I Walker, Giulio Tononi, and Larissa Albantakis. How causal analysis can reveal autonomy in models of biological systems. Philosophical transactions. Series A, Mathematical, physical, and engineering sciences, 375(2109):20160358, ec 2017.

[21] William G.P. Mayner, William Marshall, Larissa Albantakis, Graham Findlay, Robert Marchman, and Giulio Tononi. PyPhi: A toolbox for integrated information theory. PLoS Computational Biology, 14(7):e1006343, jul 2018.

[22] Brian Odegaard, Robert Knight, and Hakwan Lau. Should a few null findings falsify prefrontal theories of conscious perception? Journal of Neuroscience, 37, 2017.

[23] Masafumi Oizumi, Larissa Albantakis, and Giulio Tononi. From the Phenomenology to the Mechanisms of Consciousness: Integrated Information Theory 3.0. PLoS Computational Biology, 10(5):e1003588, may 2014.

[24] J Pearl. Causality: models, reasoning and inference, volume 29. Cambridge Univ Press, 2000.

[25] Hans Reichenbach and Maria Reichenbach. The Direction of Time. Dover Books on Physics. Dover, Mineola, N.Y, 1999.

[26] Wesley C. Salmon. Statistical Explanation and Statistical Relevance. University of Pittsburgh Press, September 1971.

[27] Giulio Tononi. An information integration theory of consciousness. BMC neuroscience, 5:42, nov 2004.

[28] Giulio Tononi. On Being. forthcoming.

[29] Giulio Tononi and Olaf Sporns. Measuring information integration. BMC neuroscience, 4(31):1–20, 2003.

[30] Peter A. White. Is conscious perception a series of discrete temporal frames? Consciousness and Cognition, 60:98–126, 2018.

